# Meta-analysis of gonadal transcriptome provides novel insights into sex change mechanism across protogynous fishes

**DOI:** 10.1101/2023.07.09.545663

**Authors:** Ryo Nozu, Mitsutaka Kadota, Masaru Nakamura, Shigehiro Kuraku, Hidemasa Bono

## Abstract

Protogyny, being capable of changing from female to male during their lifetime, is prevalent in 20 families of teleosts but is believed to have evolved within specific evolutionary lineages. Therefore, shared regulatory factors governing the sex change process are expected to be conserved across protogynous fishes. However, a comprehensive understanding of this mechanism remains elusive. To identify these factors, we conducted a meta-analysis using gonadal transcriptome data from seven species. We curated data pairs of ovarian tissue and transitional gonad, and employed ratios of expression level as a unified criterion for differential expression, enabling a meta-analysis across species. Our approach revealed that classical sex change-related genes exhibited differential expression levels between the ovary and transitional gonads, consistent with previous reports. These results validate our methodology’s robustness. Additionally, we identified novel genes not previously linked to gonadal sex change in fish. Notably, changes in the expression levels of acetoacetyl-CoA synthetase and apolipoprotein Eb, which are involved in cholesterol synthesis and transport, respectively, suggest that the levels of cholesterol, a precursor of steroid hormones crucial for sex change, are decreased upon sex change onset in the gonads. This implies a potential universal influence of cholesterol dynamics on gonadal transformation in protogyny.

## Introduction

Sequential hermaphroditism, a phenomenon in which individuals undergo sex change during their lifetime, has been extensively recognized in teleosts over the past several decades (Atz, 1964). Sequential hermaphroditism has been categorized into three major types based on the direction of sex change (protogyny, where females change to males; protandry, where males change to females; and bidirectional sex change, where individuals can change both sexes serially). Phylogenetic analysis has shown that sequential hermaphroditism has been acquired in various evolutionary lineages (Mank et al., 2006) (Fig. 1). Interestingly, protogyny is prevalent in 20 families but is thought to have emerged once in evolutionally the Percomorpha (Kuwamura et al., 2020). Therefore, there are common features in the intricate regulatory mechanisms during protogynous sex change. Specifically, the replacement of ovarian tissue with testicular tissue is commonly observed during protogynous sex change (Bhandari et al., 2003; Lo Nostro et al., 2003; Muncaster et al., 2013; Nakamura et al., 1989; Ohta et al., 2008). Additionally, considering the high functional conservation of gonads across organisms, the existence of a regulatory mechanism governing gonadal transformation comprising common and essential factors across protogynous fishes is plausible; however, a comprehensive perspective of this process remains elusive.

**Fig. 1.**
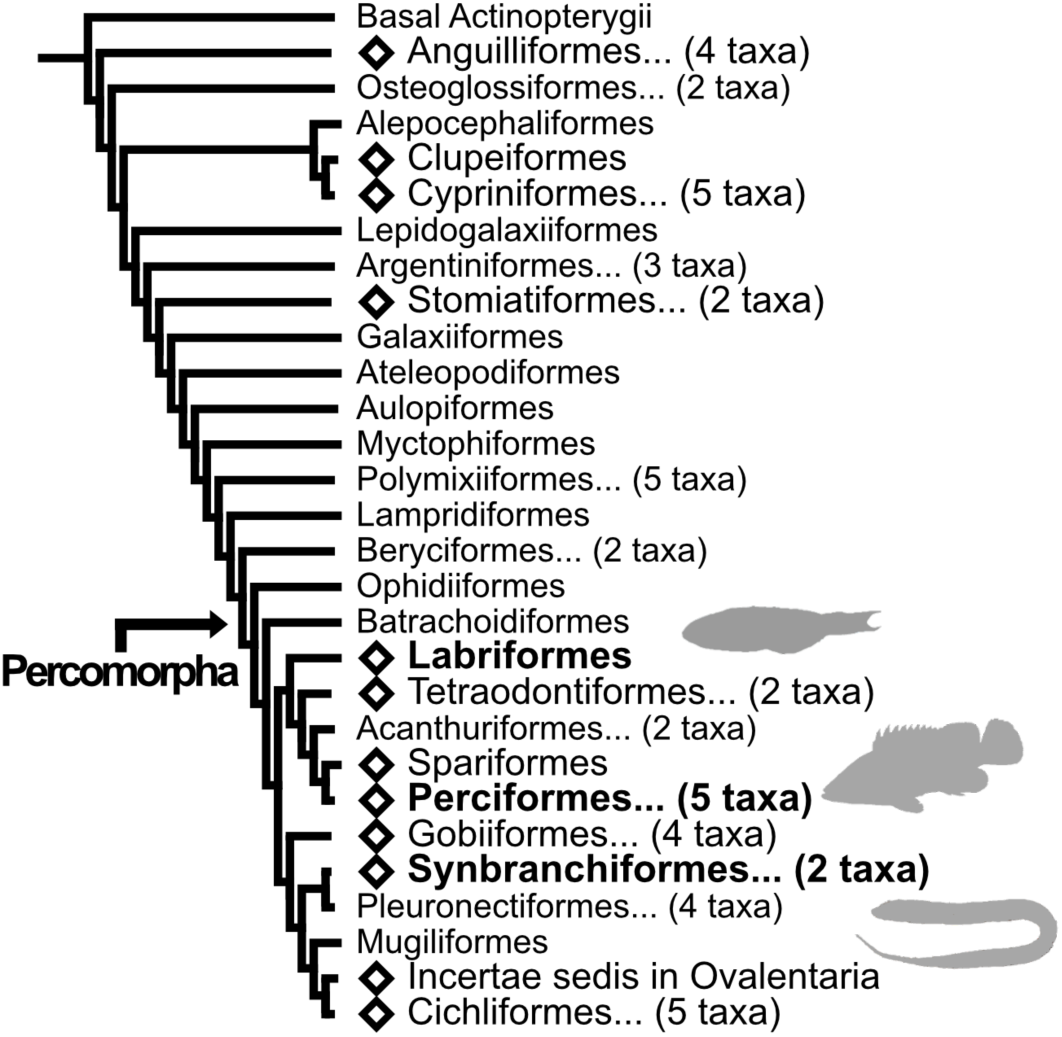
Summarized phylogenetic tree of Actinopterygii. The order-level phylogenetic tree is shown. The phylogenetic tree was derived from Rabosky et al., (2018) and modified. Diamonds indicate orders with sequential hermaphroditism. Numbers in parentheses indicate the number of summarized orders. Orders containing the fish species used in this study are shown in bold.

Accordingly, elucidation of the molecular mechanisms underlying the gametogenesis switch in sex-changing fish has subsequently become a focus of research. In particular, the expression patterns of multiple sex-related genes during sex change within the family Labridae, which serves as a model for protogynous sex change, have been explored (Horiguchi et al., 2018; Miyake et al., 2012; Thomas et al., 2019). Recently, next-generation sequencing technology has enabled researchers to conduct comprehensive analyses of gene expression, allowing the identification of global gene expression profiles during gonadal transformation in protogynous fish belonging to Labriformes, Perciformes, and Synbranchiformes (Fan et al., 2022; He et al., 2022; Senthilkumaran et al., 2017; Todd et al., 2019; Wu et al., 2020). Thus, these results suggest the involvement of several pivotal genes in the ovary-to-testis transformation in sex-changing fish.

For example, a repertoire of sex-related genes, including cytochrome P450, family 19, subfamily A, polypeptide 1a (*cyp19a1a*), forkhead box L2a (*foxl2a*), folliculogenesis specific bHLH transcription factor (*figla*), anti-Müllerian hormone (*amh*), gonadal somatic cell derived factor (*gsdf*), doublesex and mab-3 related transcription factor 1 (*dmrt1*) were identified as differentially expressed genes (DEGs) during sex change across protogynous fishes. Notably, the gonadal expression of *cyp19a1a* and *foxl2a*, which are key players in maintaining ovarian status, is reduced upon the onset of sex change in multiple species (Li et al., 2006; Thomas et al., 2019; Zhang et al., 2013). *cyp19a1a* encodes aromatase, an enzyme responsible for the conversion of androgens (e.g., testosterone) to estrogens (e.g., estradiol-17ß; E2), whereas *foxl2a* encodes a transcription factor that targets and up-regulates *cyp19a1a* in teleost fishes (Wang et al., 2007; Zhang et al., 2022). Furthermore, the initiation of female-to-male sex change, triggered by a rapid decline in E2 levels (Nakamura et al., 1989), is well-coordinated with the expression profiles of these genes. Conversely, *amh*, *gsdf*, and *dmrt1* are crucial for testicular differentiation and development in teleost fishes (Kobayashi et al., 2008; Rodríguez-Marí et al., 2005; Shibata et al., 2010; Skaar et al., 2011; Wang & Orban, 2007; Webster et al., 2017; Zhang et al., 2016). In several sex-changing species, the upregulation of these genes coincides with the emergence of testicular tissue during transition, suggesting that these genes are involved in testicular formation during gonadal sex change (Horiguchi et al., 2013; Horiguchi et al., 2018; Nozu et al., 2015; Thomas et al., 2019; Zhu et al., 2016).

Transcriptome analysis during gonadal transformation has also revealed differential expression of several genes not previously implicated in sex determination or differentiation pathways. In particular, upregulation of apolipoprotein Eb (*apoeb*), ectonucleotide pyrophosphatase/phosphodiesterase 2 (*enpp2*), and l-rhamnose-binding lectin CSL2 (*csl2*) expression, accompanied by downregulation of galectin-3 (*gal3*), was observed during the early phase of gonadal sex change in the protogynous Asian swamp eel *Monopterus albus*. These findings suggest that these genes are involved in gonadal transformation, particularly during the initiation of sex change (Fan et al., 2022; Senthilkumaran et al., 2017). However, verifying their direct involvement and commonality in the sex change process remains challenging.

Recently, by amalgamating transcriptome data from multiple studies, meta-analysis has attracted attention as a powerful tool capable of providing novel insights overlooked by traditional hypothesis-driven research methods (Bono, 2021; Rahman et al., 2020; Toga & Bono, 2023). In other words, it is expected that by reanalyzing studies across multiple species and studies, universal factors controlling gonadal transformation, which have not been detected in individual studies, could be identified among protogynous fishes. Therefore, the objective of the present study was to identify the shared factors involved in gonadal transformation across species using a meta-analysis of publicly available transcriptome data from multiple experiments and species. Furthermore, in this study, we focused on another well-studied species, the three-spot wrasse, *Halichoeres trimaculatus*, a protogynous species belonging to the family Labridae and produced *de novo* genome assembly and gonad transcriptome data undergoing sex change. This new data consolidated our dataset encompassing seven protogynous species from three orders allowing exhaustive transcriptome analysis. Through this dataset and uniform pipeline, we identified genes with differential expression levels between the transitional gonads and ovaries (Fig. S1). We reevaluated the relationship between these genes and gonadal sex change, providing a novel perspective regarding the mechanisms underlying this process.

## Results

### *De novo* genome assembly and transcriptome of gonadal tissue in the three-spot wrasse

The genome sequencing was required for the RNA-Seq analysis in this study, so the *de novo* genome assembly of the three-spot wrasse, which had not been previously determined, was constructed. High-throughput genome sequence data were obtained from a locally obtained female three-spot wrasse from Okinawa, Japan. The obtained HiFi data, which comprised approximately 32 Gbp and encompassed approximately 1.9 million DNA sequences, were assembled using the hifiasm program (Cheng et al., 2021). Subsequently, the acquired Hi-C data were used for scaffolding, resulting in the generation of 227 scaffolds. The longest scaffold measured 44.2 Mbp, with an N50 length (the shortest of the lengths covering 50% of the genome) of 37.9 Mbp, contributing to a total assembly length of approximately 849.9 Mbp. Assessment using BUSCO v5.4 (Manni et al., 2021) revealed a completeness > 98%, indicating a high level of coverage and contiguity in the *de novo* genome assembly of the three-spot wrasse (Table 1).

**Table 1.** Statistics of genome assembly in Three-spot wrasse (fHalTri1.1) and other Labidae fishes listed in NCBI Genomes.

|  | fHalTri1.1 | *Cheilinus undulatus*<br>(humphead wrasse) | *Notolabrus celidotus*<br>(New Zealand spotty) | *Labrus bergylta*<br>(ballan wrasse) | *Semicossyphus pulcher*<br>(California sheephead) | *Tautoglabrus adspersus*<br>(cunner) |
| --- | --- | --- | --- | --- | --- | --- |
| Assembly accession | BSYP01000001-BS<br>YP01000227 | GCF_018320785.1 | GCF_009762535.1 | GCF_900080235.1 | GCA_022749685.1 | GCA_020745685.1 |
| Total seq length (Mb) | 849.9 | 1173.4 | 846.7 | 805.5 | 794.1 | 723.6 |
| Max length (Mb) | 44.2 | 59.6 | 42.4 | 9.1 | 38.5 | 37.1 |
| Num of scaffolds | 227 | 45 | 467 | 13,466 | 178 | 347 |
| Scaffold N50 (Mb) | 37.9 | 51.5 | 37.1 | 0.8 | 32.1 | 31.7 |
| Scaffold L50 | 11 | 11 | 11 | 252 | 12 | 11 |
| Num of seqs > 1 Mb | 25 | 24 | 24 | 197 | 35 | 24 |
| % of seqs > 1 Mb | 97.3 | 99.7 | 97.6 | 43.9 | 97.1 | 99.9 |
| Num of gaps | 117 | 284 | 1,133 | 257 | 8 | 283 |
| Complete BUSCO (%) | 98.7* | 99.0 | 97.1 | 98.9 | 98.9 | 98.1 |

\* BUSCO version 5.4, reference datasets: vertebrata\_odb10

The RNA-Seq reads obtained from transitional gonads and ovaries of additional females were mapped to the genome assembly, and the obtained mapping results were used to assemble the transcriptome for the annotation file (“merged.gtf”). This process yielded 27,573 transcript sequences (Table 2). Subsequently, 24,695 coding sequences (protein sequences) were predicted from the set of transcript sequences using TransDecoder (Haas, 2018).

**Table 2.** Statistics of orthology relationship among species.

| Common name | Scientific name | Order | Reference genome | *Num of transcripts seqs<br>obtained based on the<br>RNA-seq data from<br>gonadal tissue | Num of protein seqs<br>predicted by<br>TransDecoder | Num of Reciprocal<br>Best Hit<br>(vs Three-spot wrasse) |
| --- | --- | --- | --- | --- | --- | --- |
| Three-spot wrasse | *Halichoeres trimaculatus* | Labriformes | Assembly in this study | 27,573 | 24,965 | 24,965 |
| Bluehead wrasse | *Thalassoma bifasciatum* | Labriformes | GCA_008086565.1 | 57,699 | 43,195 | 13,871 |
| Hong Kong grouper | *Epinephelus akaara* | Perciformes | Ge et al., 2019 | 26,507 | 24,752 | 11,619 |
| Orange-spotted grouper | *Epinephelus coioides* | Perciformes | GCA_900536245.1 | 30,867 | 34,191 | 12,507 |
| Black seabass | *Centropristis striata* | Perciformes | NA | ** 95,941 | 95,941 | 12,591 |
| Asian swamp eel | *Monopterus albus* | Synbranchiiformes | GCF_001952655.1 | 53,472 | 50,693 | 12,765 |
| Zig-zag eel | *Mastacembelus armatus* | Synbranchiiformes | GCF_900324485.2 | 48,724 | 40,908 | 13,095 |

\* Based on the annotation file (merged gtf file), the extracted transcripts sequences from the
genome by gffread
\*\* Among transcriptome sequences (594,726 seqs) assembled by Trinity, transcripts sequences
corresponding to coding sequences predicted via TransDecoder

### Data curation of public RNA-Seq data and genomic information

We downloaded Sequence Read Archive (SRA) data from six fish species for which sex changes have been reported (Kuwamura et al., 2020): bluehead wrasse *Thalassoma bifasciatum* (Todd et al., 2019) (order Labriformes), orange-spotted grouper *Epinephelus coioides* (Xiao et al., 2020) (order Perciformes), Hong Kong grouper *E. akaara* (Ge et al., 2019) (order Perciformes), black seabass *Centropristis striata* (Breton et al., 2019) (order Perciformes); Asian swamp eel *Monopterus albus* (Fan et al., 2022; Senthilkumaran et al., 2017; Zhao et al., 2018) (order Synbranchiformes), zig-zag eel *Mastacembelus armatus* (Xue et al., 2021) (order Synbranchiformes). Furthermore, using the metadata associated with the SRA data, we curated the RNA-Seq data into comparable sample pairs of intersexual (transitional) gonads and ovaries (a total of 31 paired data, including three-spot wrasse). Table S1 presents the metadata, including Run accession numbers, for the RNA-Seq data analyzed in this study.

The genome sequences of five of the seven species utilized in this study are publicly available (Table 2) and served as reference genomes for read mapping. The genome sequences of *T. bifasciatum* (GCA_008086565.1), *E. coioides* (GCA_900536245.1), *M. albus* (GCF_001952655.1), and *M. armatus* (GCF_900324485.2) were retrieved from the National Center for Biotechnology Information (NCBI) Genome Database (accessed in October 2022). We also referenced the genome information of *E. akaara* reported by Ge et al., 2019.

### Transcriptome assembly and inference of ortholog relationships

For the five species with public reference genomes, the number of transcript sequences obtained using StringTie (Pertea et al., 2015) with the “--merge” option is shown in Table 2. For black sea bass, (without a reference genome), the number of contiguous sequences assembled using Trinity (Grabherr et al., 2011) was 594,726. Among these, 95,941 transcript sequences corresponding to the predicted coding sequences (CDS) were specified as references for expression quantification using Salmon (Patro et al., 2017). The number of protein sequences predicted using TransDecoder is shown in Table 2. Orthology inference and consolidation of orthologs within the three-spot wrasse identified 7,289 orthologs that were common to all species used in this study (Table 2), and thus were considered as candidate sex-change genes.

### Identification of differentially expressed genes

The total intersexual gonad and ovary score (IO-score) was calculated for these 7,289 genes (Fig. 2a, Table S2, IO-score calculation; see Methods). Among these, 99 genes were upregulated (Fig. 2b, Table S3) and 69 genes were downregulated (Fig. 2c, Table S4). Whereas the total IO-score of these genes was occasionally derived from a single species (up: *cep97*, *zmat2*, *tmem237a*, *supt5h*; down: *znf16l*, *lrrc8db*, *sh2b1*, *tmem263*, *cyldb*, *manea*, *nol11*), in most cases, the total IO-scores were based on contributions from two or more species. Notably, the upregulated genes included *amh* and *gsdf*, which have been previously reported to play important roles in gonadal sex change, particularly in testicular formation and development, in some protogynous fish. Conversely, the downregulated genes included *cyp19a1a*, *hsd17b1*, and *foxl2a*, which are crucial for the onset of sex change (Table 3).

**Fig. 2.**
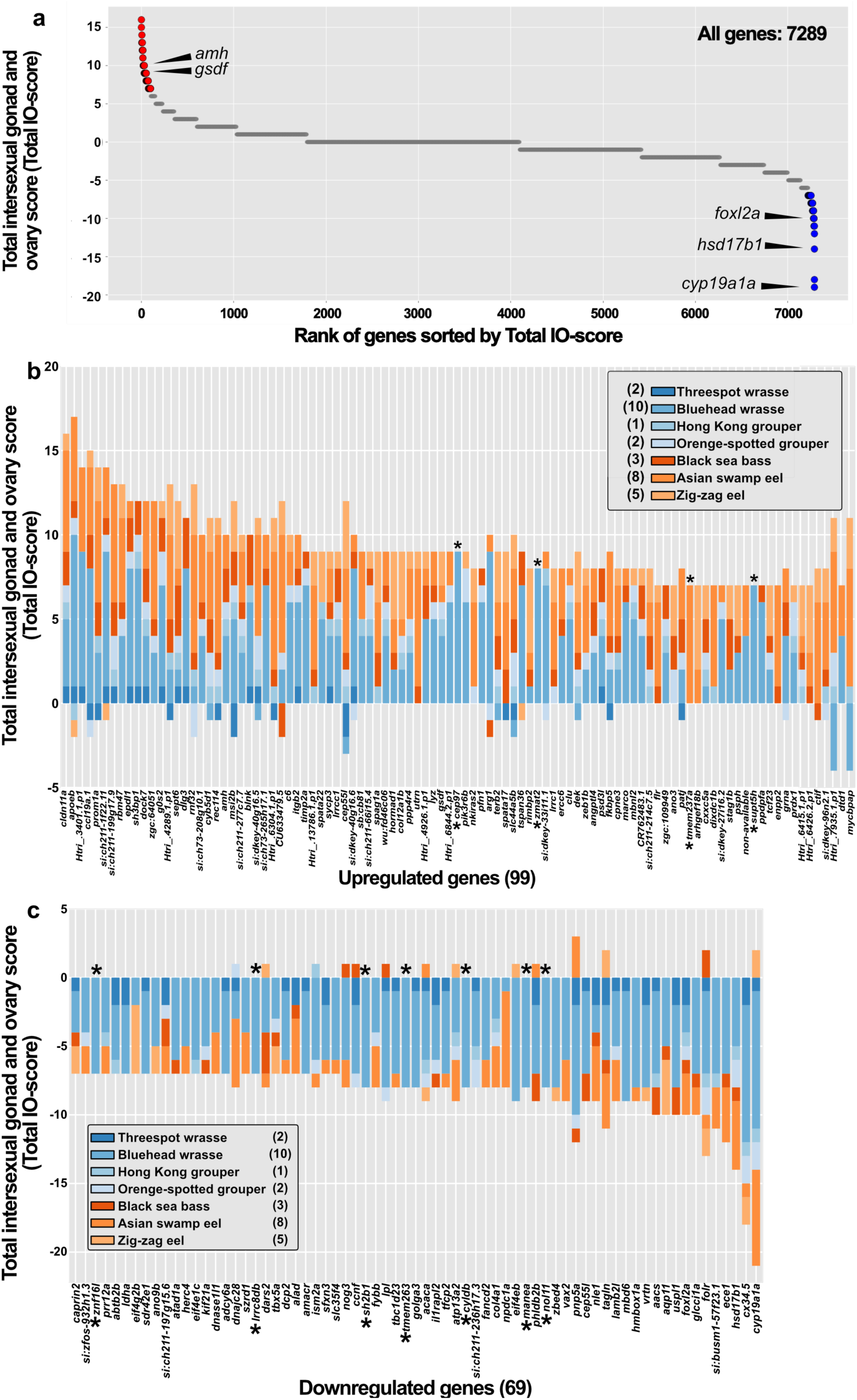
Scatter plot of the total Intersexual gonad and Ovary (IO)-score and Composition of total IO-scores for up- and downregulated genes. **a** Scatterplot of total IO-scores for the 7,289 genes in descending order. Red circles indicate 99 upregulated genes; blue circles indicate 69 downregulated genes. Gray circles (which have the appearance of lines because they overlap each other) indicate unchanged genes (6 ≤ total IO-score ≤ −6). Composition of total IO-scores for up- (**b**, n=99) and downregulated genes (**c**, n=69). Numbers in parentheses next to species name indicate the number of RNA-Seq data pairs for each species used in this study. Asterisk indicates a gene for which the total IO-score is derived from a single species.

**Table 3.**
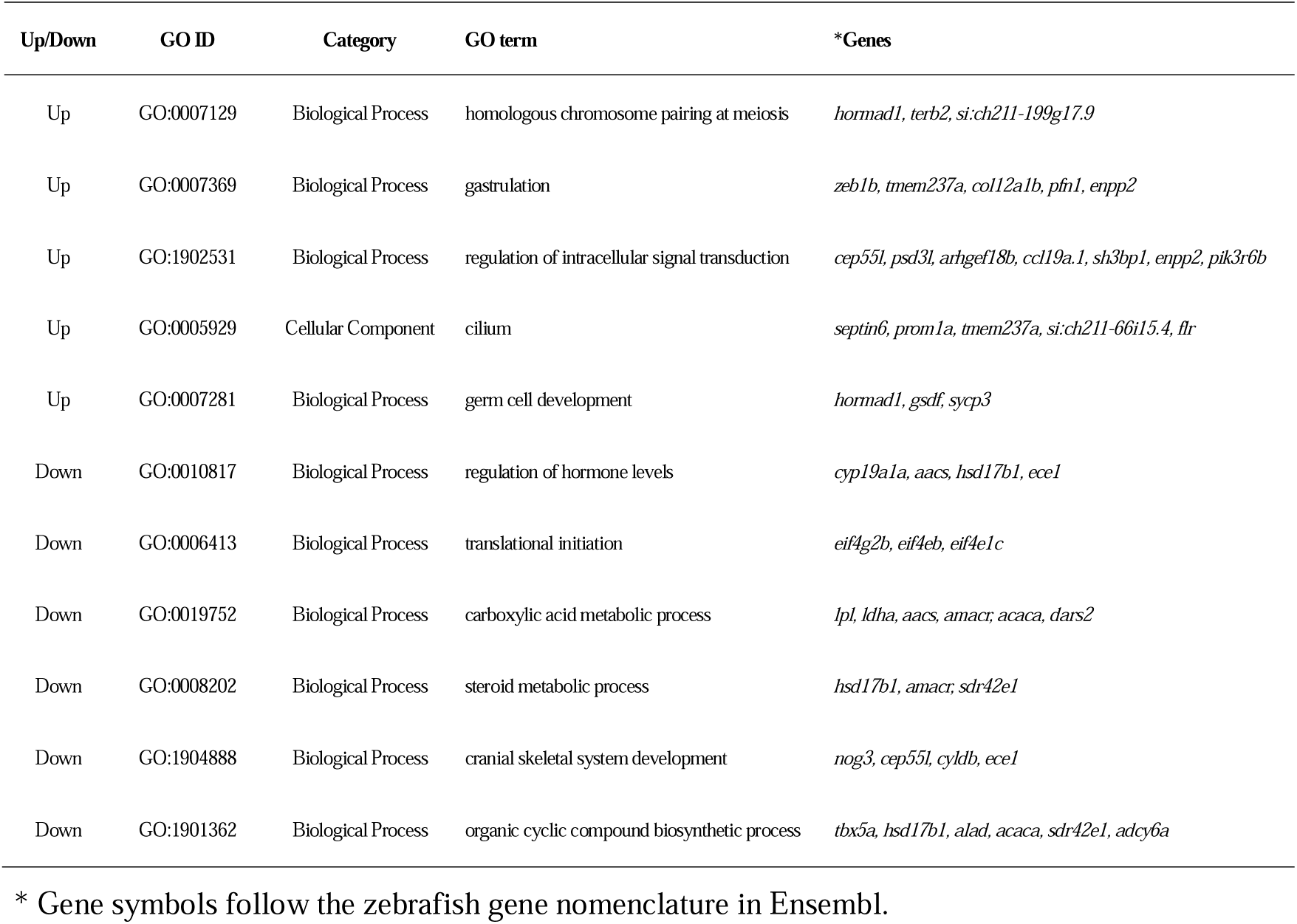
List of genes that contributed to GO term enrichment.

### Enrichment analysis of differentially expressed genes

Sequence homology search of the three-spot wrasse proteins using ggsearch36 (Pearson & Lipman, 1988) against the reference zebrafish proteins (Ensembl Release 108) (Cunningham et al., 2022) resulted in the identification of 90 homologs among the 99 upregulated genes (Table S3). No corresponding zebrafish orthologs were found for the remaining nine genes that were common to all species analyzed in this study. The results of enrichment analysis using the 90 genes that could be converted to zebrafish orthologs are shown in Fig. 3a. The Gene Ontology (GO) terms “homologous chromosome pairing at meiosis” and “germ cell development,” which are related to reproduction, were significantly enriched among the upregulated genes. The specific genes annotated with these GO terms included *hormad1*, *terb2*, *si:ch211-199g17.9*, *gsdf*, and *sycp3* (Table 3).

**Fig. 3.**
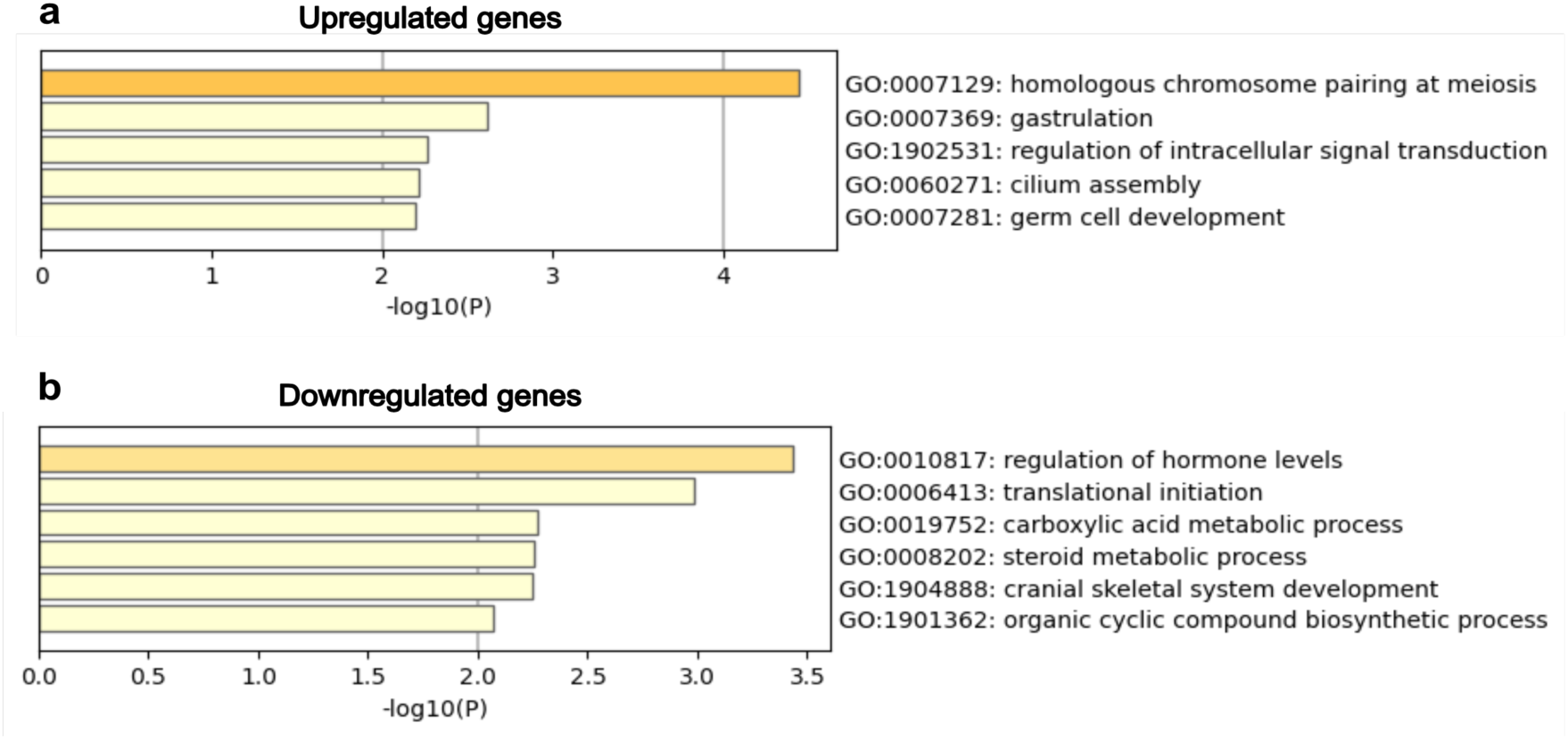
Results of enrichment analysis for differentially expressed genes using Metascape. **a** 99 upregulated genes. **b** 69 downregulated genes.

We successfully obtained the corresponding zebrafish homologs for all of the 69 downregulated genes. The results of enrichment analysis are shown in Fig. 3b. Among the downregulated genes, GO terms related to the regulation of steroid hormone levels, including “regulation of hormone levels” and “steroid metabolic process,” were significantly enriched. Specific genes annotated with these enriched GO terms included *cyp19a1a*, *aacs*, *hsd17b1*, *ece1*, *amacr*, and *sdr42e1* (Table 3).

### Pathways mapping of candidate sex-change genes

Visualization of cholesterol and steroid hormone biosynthesis pathways was conducted based on the GO terms obtained from the enrichment analysis. By referencing pathway information for teleost fishes, pathway diagrams were constructed and loaded with 7,289 genes converted to Ensembl zebrafish gene IDs (see Methods). The results showed the mapping of most enzymes involved in the pathway from acetyl-CoA to cholesterol synthesis (Fig. 4). Similarly, when mapping the steroid hormone synthesis pathway, it was confirmed that enzymes involved in the synthesis pathway from cholesterol to estradiol were included among the candidate sex-change genes (Fig. S2).

**Fig. 4.**
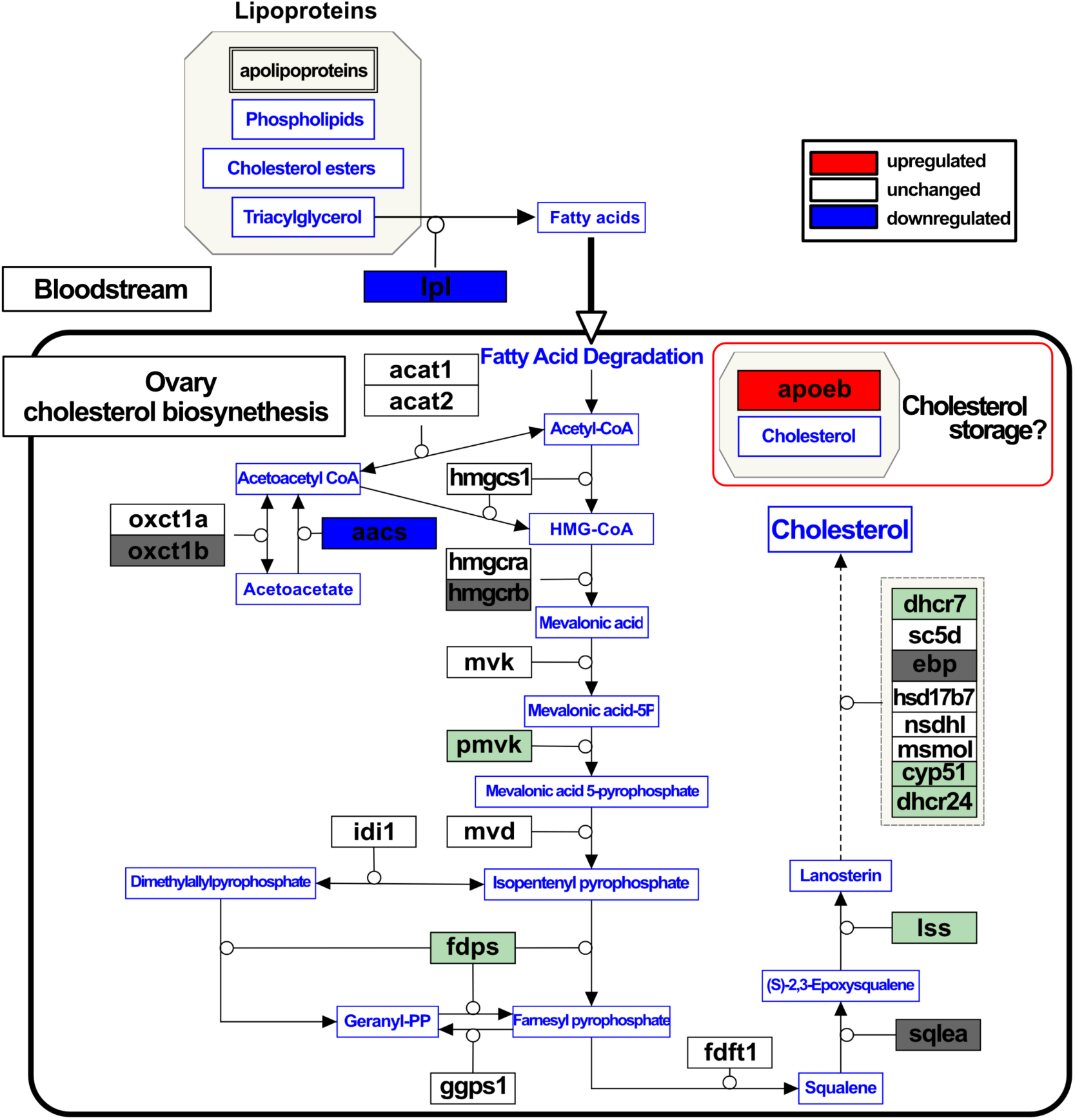
Hypothetical ovarian cholesterol biosynthesis pathway in protogynous fishes. The pathway was visualized using PathVisio (https://pathvisio.org), referencing several fish cholesterol synthesis pathways. Gene products are represented in black text, while metabolites are in blue text. Gene product annotations follow Ensembl Zebrafish gene symbols. Red-filled boxes represent gene products with confirmed upregulated expression in this study, while blue-filled boxes denote gene products with downregulated expression. Open boxes represent gene products with no expression variation. Light green-filled boxes represent gene products not included among the 7289 genes but confirmed to be present in at least the three-spot wrasse transcriptome. Dark gray-filled boxes indicate gene products that could not be identified within the three-spot wrasse transcriptome. Circles extending from gene products represent catalytic reactions, while black arrows indicate the conversion of metabolites. The dashed arrows indicate indirect pathways summarizing multi-step processes.

## Discussion

In this study, we conducted a meta-analysis of gonadal RNA-Seq data from seven protogynous fish species using a unified analysis pipeline and threshold. The aim was to compare the transcriptomes of ovaries and sexual transition gonads, with the goal of identifying common genes involved in protogynous gonadal sex change. Toward this end, we first assembled the *de novo* genome of the three-spot wrasse. The assembly statistics of the three-spot wrasse (Table 1) were comparable to those of other Labridae species, such as humphead wrasse (*Cheilinus undulatus*), ballan wrasse (*Labrus bergylta*), and New Zealand spotty (*Notolabrus celidotus*), previously deposited in the NCBI RefSeq database. These results indicated that the *de novo* genome assembly of the three-spot wrasse exhibited high quality, enabling us to perform expression analysis and orthology inference with confidence.

In our meta-analysis, we broadly examined the gonadal sex change process in protogynous fish, focusing on the replacement of ovarian tissue with testicular tissue. At a more detailed level, histological characteristics during the sex change process vary among species. In Labriformes, the early stages of sex change are predominantly characterized by ovarian tissue regression (Nozu et al., 2009; Todd et al., 2019). In contrast, in Perciformes and Synbranchiformes, testicular tissue emerges, and both ovarian and testicular tissues are present simultaneously (Liu & de Mitcheson, 2009; Senthilkumaran et al., 2017; Xue et al., 2021).

Additionally, the methods used to obtain transitional gonad samples for RNA-seq also varied in each project (Table S1). These differences are expected to impact gene expression levels. In this context, we consider the differentially expressed genes identified through the meta-analysis as crucial factors involved in gonadal transformation across species.

Through this meta-analysis, we successfully identified sex-related genes including *cyp19a1a*, *foxl2a*, *hsd17b1*, *amh*, and *gsdf*, that had been previously reported to play a role in sex change in various protogynous fish species (Fan et al., 2022; Horiguchi et al., 2013; Thomas et al., 2019; Todd et al., 2019). Furthermore, our enrichment analysis of downregulated genes highlighted a term related to the “regulation of steroid hormone production.” This finding connects to those of previous reports, which indicate that a rapid decrease in estrogen levels is crucial for female-to-male sex change (Kroon & Liley, 2000; Nakamura et al., 1989; Nozu et al., 2009). Thus, by pooling data across species, we were able to identify common genes involved in gonadal sex change. Notably, our meta-analysis demonstrates the potential of utilizing transcriptome data from multiple experiments to gain insights regarding the mechanisms underlying sex change in fish.

On the other hand, some sex-related genes, generally identified as related to sex change, showed no differential expression in this study, such as *figla*, *dmrt1*, *csl2*, and *gal3*. Particularly, *dmrt1*, *csl2*, and *gal3* were not included in the set of 7289 orthologs common to all seven species (Table S2). Among these genes, *dmrt1* is known as a key gene in male differentiation (Herpin & Schartl, 2011), and its expression profiles have also been investigated in sex-changing fish. In some protogynous fish, *dmrt1* expression increases during sex change, coinciding with the appearance of testicular tissue (Nozu et al., 2015; Todd et al., 2019). Since our dataset targets the early stages of sex change in gonads, it is suggested that the expression levels of *dmrt1* were too low for the *dmrt1* transcripts to be assembled.

Our findings demonstrate that *aacs*, annotated with the GO term “regulation of hormone levels,” is downregulated during sex change. Notably, no previous reports have suggested the involvement of *aacs* in sex change, indicating that the encoded protein (acetoacetyl-CoA synthetase; Aacs) may represent a novel gene associated with this process. Typically, AACS plays a critical role in cholesterol supply. Studies in mice have shown that *Aacs* knockdown decreases total serum cholesterol levels (Hasegawa et al., 2012). Additionally, in rat adrenal glands, inhibition of *AACS* expression through hypermethylation suppresses cholesterol supply and steroid synthesis (Wu et al., 2016). Moreover, apolipoprotein Eb (encoded by *apoeb*, listed among the upregulated genes), has been suggested to participate in cholesterol transportation in zebrafish (Gaudet et al., 2011). Apolipoprotein E, encoded by *APOE* which is the human ortholog of *apoeb*, serves as a crucial lipid transport protein involved in the modulation of cholesterol metabolism and is also synthesized within peripheral tissues (Blue et al., 1983). In addition, a previous study indicates that APOE gene expression exerts suppressive effects on steroid production in adrenal cells (Reyland et al., 1991). Specifically, the high expression of APOE in tissues with active steroid production, including the adrenal glands, appears to shift cholesterol metabolism from delivery to the steroidogenic pathway to storage of esterified cholesterol, resulting in a decrease in steroid production levels (Thorngate et al., 2002). These observations strongly support the hypothesis that the downregulation of *aacs* and the upregulation of *apoeb* during sex change contribute to suppression of the cholesterol supply and entire steroidogenesis.

Cholesterol serves as a precursor for steroid hormones, including estrogens, androgens, and glucocorticoids (Miller, 2008; Payne & Hales, 2004; Pikuleva, 2006), which play crucial roles in protogynous sex change. Previous studies have shown that a decline in endogenous estrogens, particularly E2, triggers the transition from ovary to testis in protogynous fish (Kroon & Liley, 2000; Nakamura et al., 1989; Nozu et al., 2009). E2 is synthesized from testosterone via an aromatase enzyme, encoded by *cyp19a1a*. Downregulation of *cyp19a1a* expression at the onset of sex change leads to reduced E2 production. Studies on *M. albus* and *T. bifasciatum* have revealed higher DNA methylation levels of the *cyp19a1a* gene upstream region in the testes than those in the ovaries, suggesting that epigenetic modifications may contribute to the suppression of *cyp19a1a* expression during gonadal sex change (Todd et al., 2019; Zhang et al., 2013). Our present findings could contribute one more “piece of the puzzle”, suggesting that the trigger of gonadal transformation may be attributed not only to the decreased E2 production by the downregulation of aromatase activity but also to the suppression of the cholesterol supply, the precursor of steroid hormones, leading to the inhibition of the entire steroid biosynthesis.

In fish, it is suggested that synthesized cholesterol affects the production of steroid hormones (Bera et al., 2020; Sharpe et al., 2006). This is also supported by reports that the exposure HMG-CoA reductase inhibitors in zebrafish leads to a decrease in steroid hormone levels (Al-Habsi et al., 2016). The cholesterol synthesis pathway (mevalonate pathway) primarily uses acetyl-CoA as a substrate, and *aacs* might be involved in the supply of this acetyl-CoA (Fig. 4). Additionally, a decrease in the expression of lipoprotein lipase (*lpl*) has been observed (Fig. 4, Table S4). Lipoprotein lipase is involved in the biosynthetic process of fatty acid which serves as one of the sources of acetyl-CoA. This suggests the possibility of a concurrent suppression of fatty acid supply to the gonads. Both results could lead to support the hypothesis that the decrease in steroid hormone production is due to a reduced supply of cholesterol caused by inhibition of cholesterol synthesis. Although this hypothesis requires further investigation and an understanding of cholesterol dynamics during sex change, additional exploration is warranted. Notably, a study on medaka (*Oryzias latipes*) subjected to starvation conditions implied a potential link between masculinization and lower cholesterol levels (Sakae et al., 2020). Specifically, female-to-male sex reversal was observed in starved medaka larvae, a gonochoristic fish, together with decreased cholesterol levels in the starved samples. However, the precise pathways underlying masculinization and cholesterol dynamics remain unknown. In general, cholesterol is synthesized not only in the liver and adrenal glands but also in the gonads (Bera et al., 2020; Gwynne & Strauss, 1982; Sharpe et al., 2006).

Therefore, gaining a comprehensive understanding of the lipid metabolic mechanisms in fish gonads, including cholesterol synthesis and transportation during sex change, may provide valuable insights regarding the mechanisms underlying gonadal transformation. To clarify this perspective, the remaining challenge lies in providing experimental evidence regarding whether it is possible to control gonadal transformation by actually regulating cholesterol levels within the body.

## Conclusions

In this study, our objective was to compare the transcriptomes of sex-transitional gonads with those of ovaries using a unified pipeline to identify common genes involved in gonadal sex change across protogynous fishes. We successfully constructed a high-quality *de novo* genome assembly applicable for genome-wide analysis in the protogynous three-spot wrasse and acquired newly generated gonadal transcriptome data. Leveraging this information, we conducted a comprehensive meta-analysis by integrating both public and newly derived gonadal transcriptome data from multiple protogynous fishes spanning three orders. Through this analysis, we identified 168 differentially expressed genes between the two gonadal states, suggesting their potential universal role in gonadal sex change across diverse taxa of protogynous fishes. Notably, we identified novel genes that were not previously associated with gonadal sex change. In particular, we identified cholesterol dynamics as an important candidate factor regulating the sex change process. Moreover, this finding highlights the potential of transcriptomics meta-analysis as a valuable approach to uncover overlooked mechanisms underlying conserved biological processes including sex change. We anticipate that further data-driven research utilizing these novel sex change-related genes will open new horizons in sex change research.

## Experimental procedures

### Animals

The adult female three-spot wrasse specimens collected in the field by local fisherman in the northern region of Okinawa, Japan were purchased and employed for genome and RNA sequencing. These specimens were subsequently cultivated in 1000 L tanks receiving a continuous supply of unregulated seawater, and were subjected to natural photoperiod conditions until sampling.

The maintenance and handling of fish in this study were conducted in accordance with the “Guidelines for Animal Experiments of the Okinawa Churashima Foundation,” with the same consideration for animal care and welfare as that for higher vertebrates, including reptiles, birds, and mammals. Animal ethical review and approval were not required for the present study because the guidelines stipulated that approval from the Institutional Animal Care and Use Committee of Okinawa Churashima Foundation is required for “higher” vertebrates, and is waived for “lower” vertebrates including fish.

### Genome sequencing and assembly for the three-spot wrasse

Muscle tissue from a mature female individual was used for genome sequencing. The specimen was randomly selected from cultivated tank. After anesthetizing the fish with 0.05% v/v 2-phenoxyethanol (FUJIFILM Wako Pure Chemical), the muscle tissue was dissected, immediately frozen in liquid nitrogen, and stored at −80 °C until genomic DNA extraction. High-molecular-weight DNA was extracted from muscle using a NucleoBond AXG column (Macherey–Nagel), followed by purification with phenol–chloroform. The concentration of the extracted DNA was measured using Qubit (Thermo Fisher Scientific Inc., Waltham, MA, USA), and its size distribution was first analyzed with a TapeStation 4200 (Agilent Technologies Inc., Santa Clara, CA, USA) to ensure high integrity, and subsequently analyzed via pulse-field gel electrophoresis on a CHEF DR-II system (Bio-Rad Laboratories, Inc., Hercules, CA, USA) to ensure a size range of 20–100 kb.

To obtain long-read sequence data, a SMRT sequence library was constructed using the SMRTbell Express Template Prep Kit 2.0 (PacBio, Menlo Park, CA, USA) and sequenced on a PacBio Sequel II system. The sequencing output was processed to generate circular consensus sequences to obtain HiFi sequence reads. Adapter sequences were removed from the HiFi reads using HiFiAdapterFilt (Sim et al., 2022). The obtained HiFi sequence reads were assembled using the hifiasm (v0.16.1) assembler (Cheng et al., 2021) with default parameters. The obtained contigs were first scaffolded with the P_RNA_scaffolder (Zhu et al., 2018) and further scaffolded using Hi-C data as follows. The Hi-C library was prepared with the *in situ* method using the muscle tissue of the individual used for DNA extraction according to the iconHi-C protocol (Kadota et al., 2020), employing the restriction enzymes DpnII and HinfI, and sequenced on a HiSeq X sequencing platform (Illumina, Inc., San Diego, CA, USA). The obtained Hi-C read pairs, amounting to 232 million read pairs, were aligned to the HiFi sequence contigs using the Juicer program (Durand et al., 2016), and the HiFi sequence contigs were scaffolded via the 3d-dna (version 201008) to be consistent with the chromatin contact profiles through manual curation on JuiceBox v1.11.08 (Dudchenko et al., 2018). The continuity and completeness of the resulting genome assembly were assessed using the webserver gVolante (Nishimura et al., 2017), in which the pipeline BUSCO (v5.4, reference datasets: vertebrata_odb10) was implemented (Manni et al., 2021).

### RNA-Seq of gonadal tissue in the three-spot wrasse

Based on previous studies (Horiguchi et al., 2013; Nozu et al., 2009), we triggered a sex change in three-spot wrasse n=5 females (body length; 8.0–9.3 cm, body weight; 12.4–20.9 g) to obtain transitional gonads through treatment with an aromatase inhibitor (AI), which exerts inhibitory effects on the enzyme responsible for estrogen synthesis. The collected gonads were cut into small pieces, one piece was immersed in TRIzol reagent (Invitrogen) and the rest was fixed in 4% paraformaldehyde/1× phosphate-buffered saline (PBS) for histological observation. Total RNA in each sample was extracted promptly in accordance with the TRIzol instruction manual. The extracted total RNA was dissolved in RNase-free water and stored at −80 °C until RNA-Seq analysis. We selected two treated and two control samples for RNA-Seq analysis on the basis of histological observation (Fig. S3).

Beads with oligo (dT) were used to isolate poly(A) mRNA after the total RNA was collected from the samples. Fragmentation buffer was added to disrupt mRNA into short fragments. Using these short fragments as templates, a random hexamer primer was used to synthesize the first-strand cDNA. Second-strand cDNA was synthesized using buffer, dNTPs, Rhase H, and DNA polymerase I. Short fragments were purified using the QiaQuick PCR extraction kit (Qiagen) and resolved with EB buffer for end reparation and adding an “A” base. Subsequently, the short fragments were connected to sequencing adapters. Following agarose gel electrophoresis, suitable fragments were selected as templates for PCR amplification.

Finally, the library was sequenced using an Illumina HiSeq 2000 as 2 × 90 bp paired-ends. The acquired reads were subjected to the quality control procedures described below and used for downstream expression analysis.

### Retrieval and acquisition of RNA-Seq data in protogynous fishes from public resources

To obtain RNA-Seq data from the gonads of protogynous fishes in public databases, we accessed SRA (Kodama et al., 2011) (accessed in October 2022) to search SRA data containing the following keywords: “bony fishes (AND ovary OR gonad).” SRA data were converted into FASTQ files using the Fasterq-dump program of the SRA Toolkit (v3.0.0).

### Quality control of RNA-Seq data

To ensure the integrity of the RNA-Seq reads, trimming and quality control procedures were performed. Specifically, we used Trim Galore (v0.6.7) with the option “--quality 20 --paired” to eliminate low-quality bases from paired-end reads.

### Read mapping against the reference genome

For species with available genomic information, RNA-Seq reads were mapped to the reference genome sequences using HISAT2 (v2.2.1) (Kim et al., 2019) with the parameter “-q -dta -x”. The resulting SAM files were sorted and converted into BAM files using SAMtools software (v1.15.1) (Danecek et al., 2021).

### Generating General Transfer Format (gtf) files based on genome sequence and RNA-Seq reads for reference gene annotation when quantifying expression values

To ensure consistency in the use of annotation information for expression quantification in species with available reference genomes, we employed the merge function of StringTie (v2.1.7) (Pertea et al., 2015) to generate a gtf file that served as a guide for annotations. The transcriptome assembly for each fish was conducted using StringTie with default settings based on the mapping results (BAM file). The resulting gtf files were then merged using StringTie with the “--merge” option, producing a unified file as the “merged.gtf”. This “merged.gtf” file contained a non-redundant set of assembled transcripts for each species. Generated “merged.gtf” files in each species were designated as the reference annotation files for quantifying transcript expression.

### Quantification of transcripts expression and detection of differentially expressed transcripts

When reanalyzing high-throughput data from multiple experiments, it is crucial to consider batch effects (Leek et al., 2010). In the present study, we, therefore, calculated the transcript per million (TPM) value for each transcript and utilized the difference in TPM values between the paired data as an indicator for identifying differentially expressed transcripts.

Prior to expression quantification, we performed a principal component analysis (PCA) using count data from mapped reads to examine the expression profiles of ovaries and gonads in each species (Fig. S4). The PCA and visualization were conducted using the ’pcaExplorer’ package (version 2.30.0) (Marini & Binder, 2019) in the R program.

Transcript expression was quantified in species with available genome annotation using StringTie, specifying the parameters as “-G merged.gtf -e”. The resulting gtf file for each sample provided the TPM values, which were used for downstream analysis.

TPM values were computed for species lacking reference genomes using Salmon (v.1.9.0) (Patro et al., 2017) with the parameters “quant -I index -l A”. To generate the transcriptome sequences required for Salmon, we performed transcriptome assembly using the Trinity (v2.13.1) assembler (Grabherr et al., 2011) with default settings. Open reading frames (ORFs) and CDS were predicted from the transcriptome assembly using TransDecoder (v5.5.0) (Haas, 2018) with the same settings as described below. The resulting gff3 file obtained from the TransDecoder run was used to extract transcript sequences corresponding to the predicted CDS from the transcriptome assembly. The extracted sequence set was then utilized as the reference transcriptome after eliminating duplicate sequences using seqkit (v2.3.0) (Zou et al., 2016) with the “rmdup -s” command.

To assess changes in transcript expression, the expression ratio was determined using TPM values obtained from intersexual gonad and ovary transcriptomes (referred to as the IO-ratio). The IO-ratio for each transcript was calculated using Equation (1).

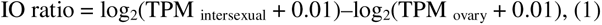

This calculation was performed for all pairs of intersexual gonad and ovary transcriptomes. To convert zeros to logarithms, a value of “0.01” was added to the TPM of each transcript.

Transcripts with an IO-ratio > log_2_(5) (indicating a 5-fold expression change) were classified as “upregulated,” whereas transcripts with an IO-ratio < −log_2_(5) (representing a 0.2-fold expression change) were considered “downregulated.” Transcripts outside these thresholds were deemed “unchanged.” The threshold was set based on the ±2 SD (standard deviation) range of expression ratios of transcripts in each species (Table S5). To determine differential expression between the intersexual gonad and ovary, an IO-score was calculated for each transcript by subtracting the count of “downregulated” pairs from that of “upregulated” pairs, as shown in Equation (2).

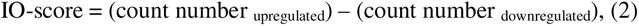

Using orthology inference as described below, the total IO-score for each transcript was calculated by summing the IO-scores of all species. Total IO-score thresholds of 7 and −7 were employed to identify transcripts that were differentially upregulated and downregulated, respectively. The thresholds were arbitrarily set based on 20% above the maximum or below the minimum values (31 or -31). All calculations for IO-ratios and IO-scores were performed using R (v 4.2.1) (R Core Team, 2022).

### Orthology inference among species

GFFread (v0.12.1) (Pertea & Pertea, 2020) was utilized along with the generated “merged.gtf” file to extract transcript sequences from the reference genomes. For species without reference genomes, transcript sequences were obtained via assembly using Trinity, as described previously. TransDecoder was used to identify the coding regions and translate them into amino acid sequences for all species. ORFs were extracted from the transcriptome sequences using TransDecoder Longorfs with the parameters “-m 100”. CDS were then predicted using TransDecoder Predict with default settings, and the protein sequences converted from the predicted CDS were acquired. Subsequently, a reciprocal homology search was conducted between the predicted protein sequences of the three-spot wrasse and those of other species, and the reciprocal best hit (RBH) was identified as the ortholog. The program Diamond (v2.0.15) (Buchfink et al., 2014) was used for the RBH search using the parameters “--ultra-sensitive -e 1e-3”. To extract the orthologs shared among all species, we consolidated the RBH results with the three-spot wrasse-predicted protein IDs.

### Gene set enrichment analysis

Differentially expressed gene sets were analyzed using the web tool Metascape (v3.5) (Zhou et al., 2019), which enables gene set enrichment analysis. However, Metascape only supports the analysis of zebrafish among teleosts. Therefore, prior to analysis, the predicted protein sequences of the three-spot wrasse were mapped to the corresponding zebrafish Ensembl protein IDs. To obtain orthologous information, we utilized the ggsearch36 program from the FASTA package (Pearson & Lipman, 1988) to globally align the predicted protein sequences of the three-spot wrasse with the proteome sequences of zebrafish (Ensembl Release 108) (Cunningham et al., 2022).

### Visualization of cholesterol and steroid hormone biosynthesis pathways and orthologs mapping to the pathways

The hypothetical cholesterol biosynthesis and steroid hormone biosynthesis pathway in teleost fish was illustrated using PathVisio (Murphy et al., 2015). Modifying information from WikiPathways (Pico et al., 2008) intended for zebrafish (WP1387, https://www.wikipathways.org/instance/WP1387) and data from the yellow perch, *Perca flavescens* (Kemski et al., 2020), facilitated the depiction of the cholesterol biosynthesis pathway. Additionally, the steroid hormone biosynthesis pathway was constructed by adjusting information sourced from various fish models (Craft et al., 2015; Fernandino et al., 2013; Li et al., 2006; Liu et al., 2022; Nagahama et al., 2021). Moreover, through PathVisio, 7,289 candidate sex-change genes, converted into Ensembl zebrafish gene IDs, were mapped onto the established biosynthesis pathways.

### Availability of data

All sequencing data, including assembled sequences and raw sequence reads, were deposited in the DNA Data Bank of Japan (DDBJ) under the umbrella BioProject accession number PRJDB15835. The genome assembly was deposited in the DDBJ under accession numbers BSYP01000001–BSYP01000227. Additionally, raw sequence reads were deposited in the DDBJ under the following accession numbers: DRR466686 (PacBio HiFi reads), DRR486910 (Illumina HiC reads), and DRR483512–DRR483515 (Illumina RNA-Seq reads).

## Competing interests

The authors have no conflicts of competing interest to declare that are relevant to the content of this article.

## Author contributions

RN and HB conceived and designed the study. MK and SK performed genome sequence and de novo genome assembly of three-spot wrasse. RN & MN RNA-seq data production of three-spot wrasse. RN performed RNA-seq data analysis and orthology inference. RN, MN, SK and HB wrote a draft manuscript. All authors read and approved the final manuscript.

## Acknowledgements

All authors thank Dr. Osamu Nishimura at DNA Sequencing Facility operated by Laboratory for Phyloinformatics in RIKEN BDR for assistance in Hi-C data assessment. We thank the Sesoko Station, Tropical Biosphere Research Center, University of the Ryukyus for allowing us to use their research facility. This research was supported by the Center of Innovation for Bio-Digital Transformation (BioDX), a program on the open innovation platform for industry-academia co-creation (COI-NEXT), and the Japan Science and Technology Agency (JST, COI-NEXT, JPMJPF2010). This work was partially received financial support by International Research Organization for Advanced Science and Technology, Kumamoto University, Kumamoto, Japan.

## Notes

### Competing Interest Statement

The authors have declared no competing interest.

### Summary of Updates

Revisions in Introduction added. Supplemental figure 5 added.

