## Supplemental Figures for "Meta-analysis of gonadal transcriptome provides novel insights into sex change mechanism across protogynous fishes"

Supplementary figure 1.

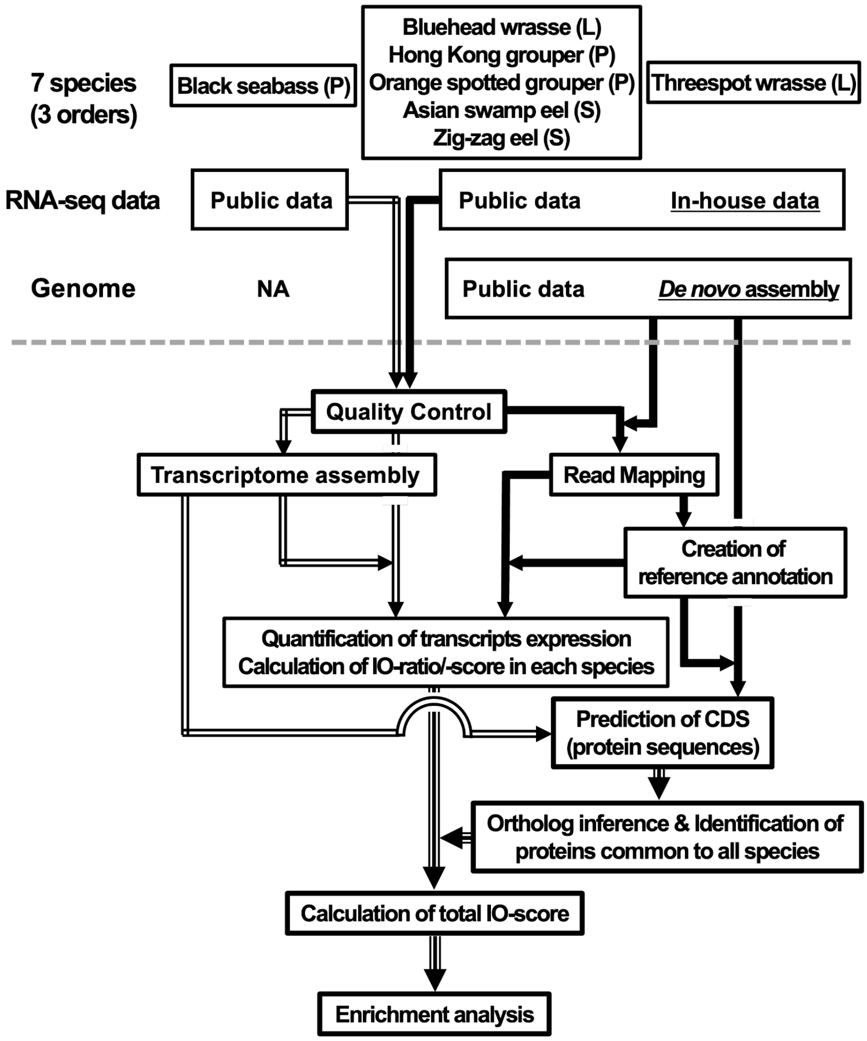


**Figure S1**. Schematic diagram of the analysis pipeline in this study. The consistent line style of the arrows indicates the utilization of common tools for conducting data processing. The letters in parentheses next to the species names represent the following order names; L: Labriformes, P: Perciformes, S: Synbranchiformes.

Supplementary figure 2.

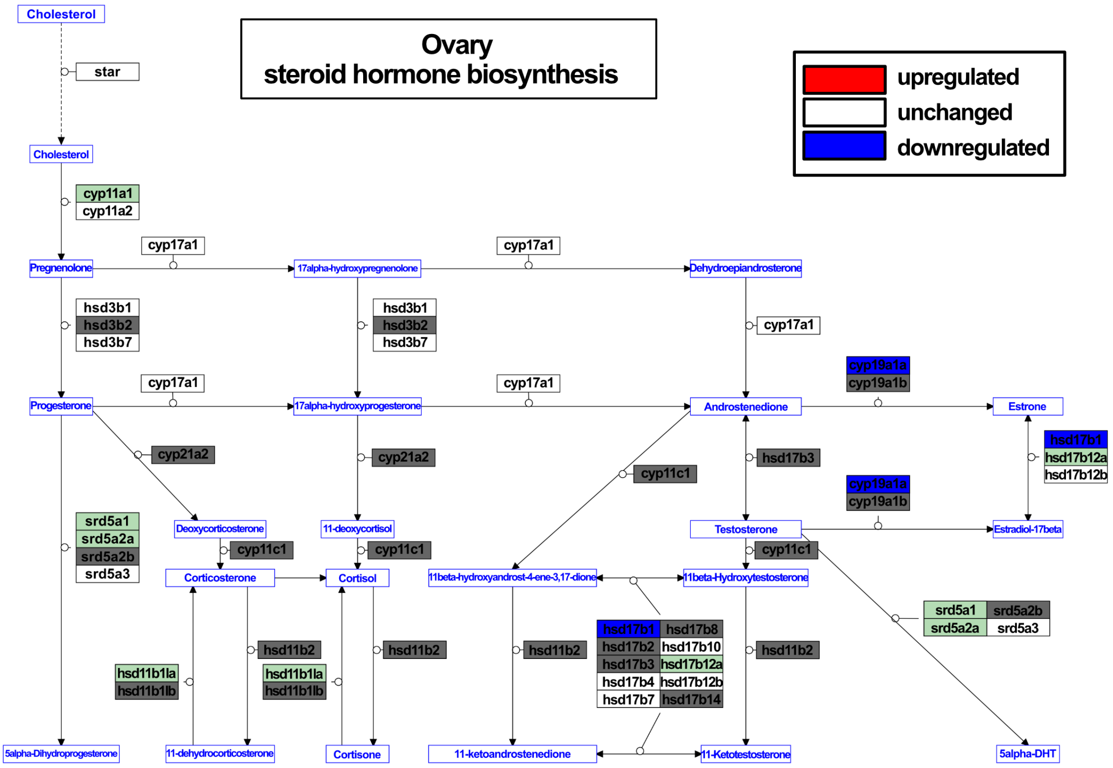


**Figure S2.** Hypothetical ovarian steroid hormones biosynthesis pathway in protogynous fishes. The pathway was visualized using PathVisio (https://pathvisio.org), referencing several fish cholesterol synthesis pathways. Gene products are represented in black text, while metabolites are in blue text. Gene product annotations follow Ensembl Zebrafish gene symbols. Red-filled boxes represent gene products with confirmed upregulated expression in this study, while blue-filled boxes denote gene products with downregulated expression. Open boxes represent gene products with no expression variation. Light green-filled boxes represent gene products not included among the 7289 factors but confirmed to be present in at least the three-spot wrasse transcriptome. Dark gray-filled boxes indicate gene products that could not be identified within the three-spot wrasse transcriptome. Circles extending from gene products represent catalytic reactions, while black arrows indicate the conversion of metabolites.

Supplementary figure 3.

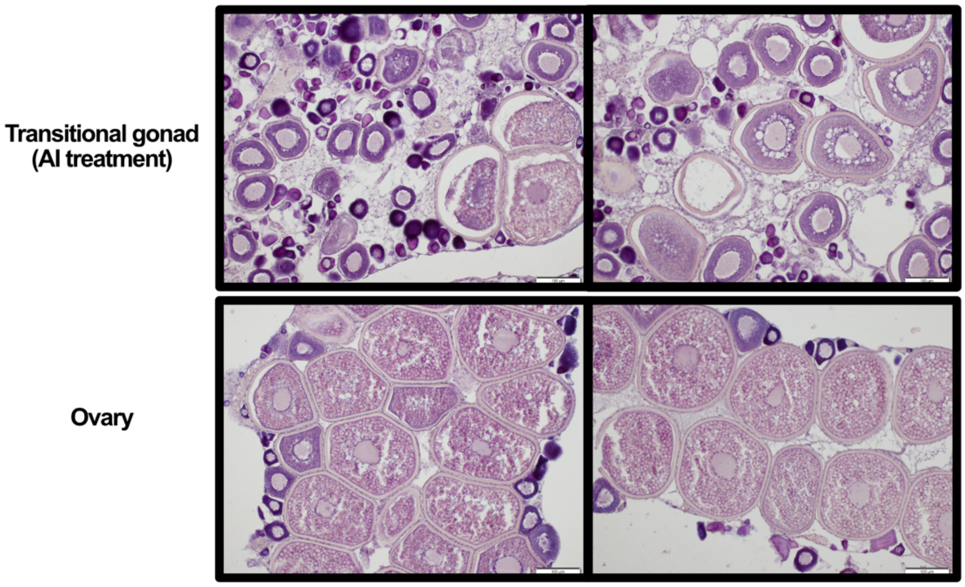


**Figure S3.** Histological sections of control ovaries and transitional gonads treated AI for 7 days in the threespot wrasse, *Halichoeres trimaculatus*.

Supplementary figure 4.

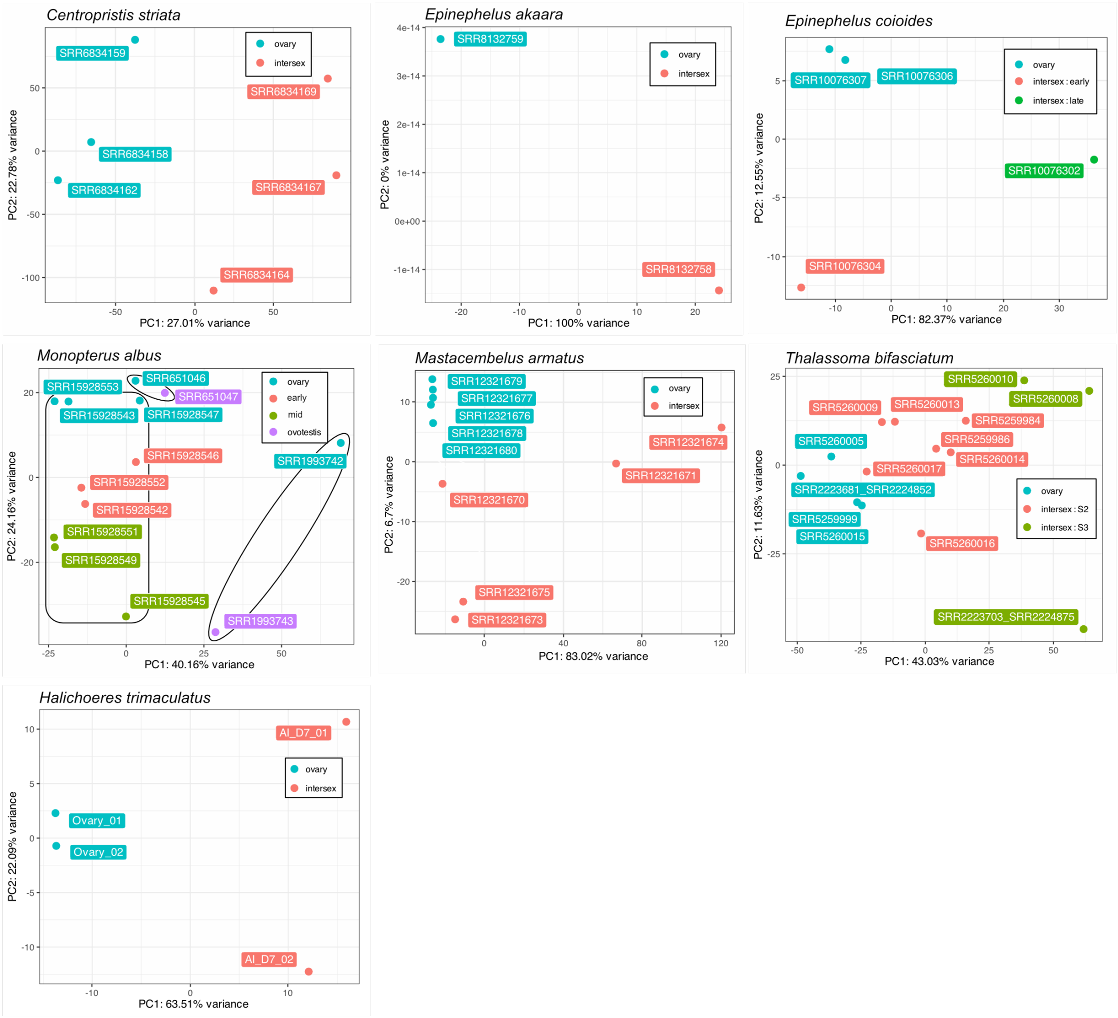


**Figure S4.** Principal component analysis (PCA) results of gene expression data from each species used in this study. The status of transitional gonads is indicated according to the metadata descriptions in the supplementary table 1. The black line circles in *Monopterus albus* represent data derived from different bioprojects. The analysis and visualization were performed using the R program (version 4.4.0) with the 'pcaExplorer' package (version 2.30.0; Marini and Binder, 2019).
